# Spatial structure increases the benefits of antibiotic production in *Streptomyces griseus*

**DOI:** 10.1101/482786

**Authors:** Sanne Westhoff, Simon B. Otto, Aram Swinkels, Bo Bode, Gilles P. van Wezel, Daniel E. Rozen

**Affiliations:** Institute of Biology, Leiden University, Sylviusweg 72, 2333 BE Leiden, The Netherlands

## Abstract

Bacteria in the soil compete for limited resources to survive and proliferate. One of the ways they might do this is by producing antibiotics, but the costs of antibiotic production and their low concentrations in soils have led to uncertainty about the role of these natural products for the bacteria that produce them. Here, we examine the fitness effects of streptomycin production by the filamentous soil bacterium *Streptomyces griseus* and the conditions that modify its ability to invade competitors. Using pairwise competion assays, we first provide direct evidence that streptomycin production enables *S. griseus* to kill and invade a population of the susceptible species, *S. coelicolor*, but not a streptomycin-resistant mutant of this species. Next we show that the fitness benefits of streptomycin production are density-dependent, because production scales positively with cell number, and frequency-dependent, with a threshold of invasion of *S. griseus* at around 1%. Finally, using serial transfer experiments where spatial structure is either maintained or periodically destroyed, we show that spatial structure reduces the threshold frequency of invasion by more than 100-fold, indicating that antibiotic production can permit invasion from extreme rarity. Our results provide clear evidence that streptomycin is both an offensive and defensive weapon that facilitates invasion into occupied habitats and also protects against invasion by competitors. They also indicate that the benefits of antibiotic production rely on ecological interactions occurring at small local scales, suggesting that low antibiotic concentrations in bulk soil are unlikely to be representative of their effective concentrations in nature.

## Introduction

Soil is a heterogeneous habitat where bacteria have to compete for limited resources to survive and proliferate. Bacteria have evolved different strategies to compete with their neighbours. They become motile in search for more favourable conditions, compete by increasing resource uptake and assimilation, or compete by interference by producing toxins, like bacteriocins or antibiotics (Ghoul & Mitri, 2016; Hibbing et al., 2010). Antibiotic production may allow producing strains to inhibit or kill their competitors, thereby allowing increased access to resources or space. However, antibiotics are costly to produce and the overall concentrations of antibiotics in the soil are low (Yim *et al.*, 2006), raising questions about their role in nature and the conditions that enable antibiotic producing bacteria to become established.

Antibiotics have been traditionally viewed as antibacterial weapons that are used defensively to prevent competitors from invading an already colonized niche, or offensively, to invade and displace competing bacteria (Raaijmakers and Mazzola, 2012; Wiener, 1996). More recently, antibiotics have been instead suggested to act as signalling molecules and regulators in microbial communities (Davies et al., 2006; Martínez, 2008). A key argument of this alternative hypothesis is that antibiotic concentrations in nature are too low to kill competitors (Davies, 2006). Three arguments highlight problems with these concerns. First, because antibiotic concentrations will vary spatially according to the distance from the producing strain, average concentrations measured in bulk soil are unlikely to be informative of “effective” concentrations at a more local scale. Second, recent experiments have clarified that even sub-inhibitory concentrations of antibiotics strongly select for drug resistance, suggesting that low levels of production may be sufficient to provide antibiotic producing bacteria with direct benefits (Gullberg et al., 2011, Westhoff et al., 2017). Finally, the widespread presence of antibiotic resistance genes in pristine environments that are unaffected by anthropogenic antibiotic contamination indicates that antibiotic production has had a pronounced effect on bacterial communities (D’Costa *et al.*, 2006; Martínez, 2008).

Although these results, together with other studies indicating an aggressive function for antibiotics, seem to favour the traditional view of antibiotics as weapons, there are few studies that directly quantify the fitness effects of antibiotic production or that clarify their role as either offensive or defensive weapons for interference competition (Wiener, 1996, 2000; Abrudan *et al.*, 2015).

Here we study the fitness effects and establishment conditions of antibiotic production in the filamentous soil-dwelling bacterial species *Streptomyces griseus*. *Streptomyces* are prolific producers of antibiotics, including more than half of the antibiotics used in clinical practice, as well as a diversity of other secondary metabolites with anti-fungal, anti-parasite or anti-cancer activities (Bérdy, 2005; Barka *et al.*, 2016; Hopwood, 2007). *S. griseus* is of both historical and ecological relevance. This species produces streptomycin, a broad-spectrum aminoglycoside antibiotic that inhibits translation and was the first clinically deployed antibiotic from Streptomyces to be discovered (Schatz *et al.*, 1944; Pfuetze *et al.*, 1955). *S. griseus* is also biogeographically widespread, with significant variation in streptomycin resistance (Laskaris *et al.*, 2010) and production (preliminary data) across natural isolates. By examining the population dynamics of competition experiments between *S. griseus* and a streptomycin-sensitive competitor, *S. coelicolor*, we provide clear evidence that streptomycin is an offensive weapon that facilitates invasion, while also showing that the capacity for invasion varies with population density and frequency and is significantly facilitated by spatial structure.

## Results

### Streptomycin enables S. griseus to invade S. coelicolor

We first asked whether *S. griseus* could invade a population of streptomycin susceptible *S.coelicolor* (MIC 2 μg/ml) by growing the two species together in paired competition. Both competitors were mixed at equal densities and a total of 10^5^ spores was plated and then, after growth, spores were harvested 4 days later. Importantly, when grown alone under these conditions, *S. coelicolor* produces significantly more spores than *S. griseus*, (Fig. S1, unpaired t-test, df = 4, p = 0.002), leading to the null hypothesis that this species would be competitively dominant. However, when the two species were mixed 1:1, *S. griseus* readily displaced *S. coelicolor* (Fig. 1A), a result that is due to the decline of the susceptible species. And when we repeated this experiment using strains of *S. coelicolor* with decreased susceptibility to streptomycin (MIC ranging from 12 – 192 μg/ml), we found that the fitness of *S. griseus* declined (Fig 1B). These results provide direct evidence that streptomycin production, rather than any other secreted metabolites of *S. griseus*, allows this species to invade *S. coelicolor*. The data also suggest that *S. griseus* at these densities produces quite high concentrations of streptomycin, because only the strain of *S. coelicolor* with the highest MIC (192 μg/ml) was able to prevent *S. griseus* invasion.

**Fig. 1.**
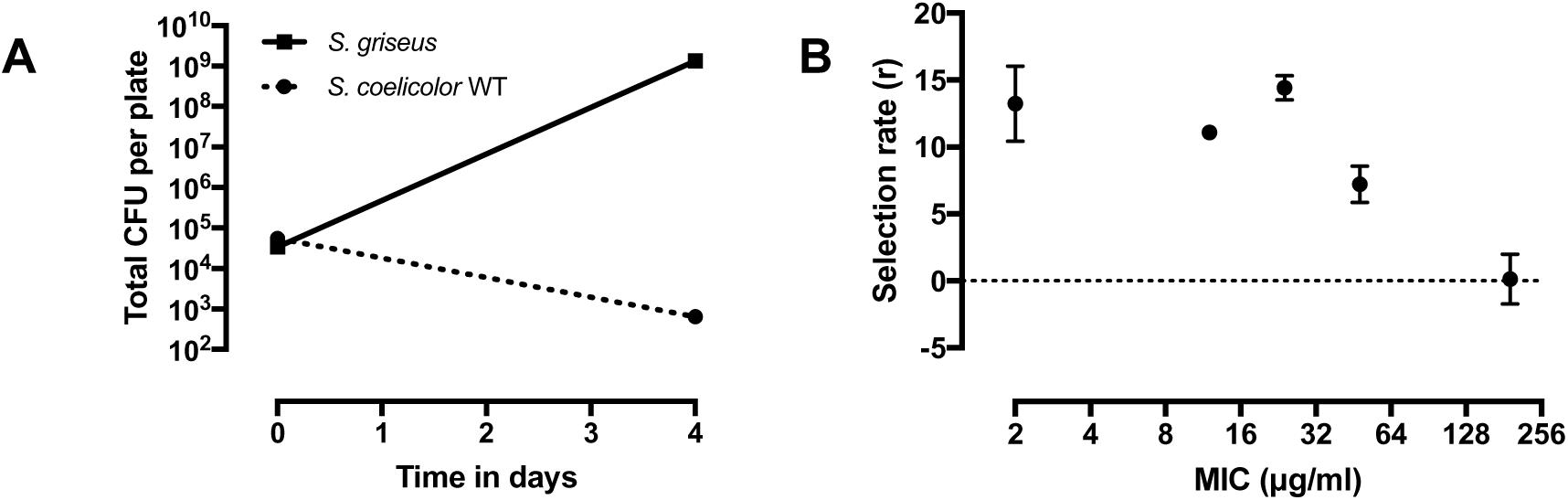
Competition between *S. griseus* and *S. coelicolor* mixed at equal frequencies. (A) Initial and final densities of *S. griseus* and a fully susceptible strain of *S. coelicolor* during 4 days of pairwise competition (B) Fitness of *S. griseus* competed against susceptible *S. coelicolor* WT (MIC 2 μg/ml streptomycin), intermediate resistant (MIC 12, 24 or 48 μg/ml) or high level resistant (MIC 192 μg/ml) *S. coelicolor* mutants. Error bars represent standard errors of the mean.

### Streptomycin production increases with inoculation density

To estimate how much streptomycin *S. griseus* produces we measured the size of inhibition zones against *S. coelicolor* and compared these to zones produced by known concentrations of purified streptomycin. We first generated a standard curve by extracting agar plugs from plates made with increasing concentrations of streptomycin and then placing these on a plate inoculated with *S. coelicolor*. As expected, this revealed that halos became larger with increasing streptomycin concentrations for plates inoculated with the WT strain. For plates inoculated with strains with intermediate levels of resistance (MIC equal to 12 and 24 μg/ml streptomycin) smaller halos appeared only at higher streptomycin concentrations, and halos were absent in the strain with the highest MIC (192 μg/ml) (Fig. 2A). These results were used to estimate streptomycin production by *S. griseus* by excising agar plugs from plates inoculated with different densities of *S. griseus* after growth for four days (Fig. 2B). As with pure streptomycin (Fig. 2A), we found that the size of the zone of inhibition increased for the susceptible strain of *S. coelicolor* but saturated at low densities of *S. griseus*, reaching a maximum halo size at an inoculum density of ~10^3^ *S. griseus* spores. This indicates that even at low densities (in monoculture) *S. griseus* produces as much antibiotic as a pure streptomycin stock prepared at 128 ug/ml. As with the pure streptomycin, we observed smaller halo sizes when *S. griseus* plugs were placed on plates containing *S. coelicolor* strains with intermediate streptomycin resistance and no halos with the high-level resistant strain (Fig. 2B). These results show that streptomycin production occurs at high levels and is density dependent.

**Fig. 2.**
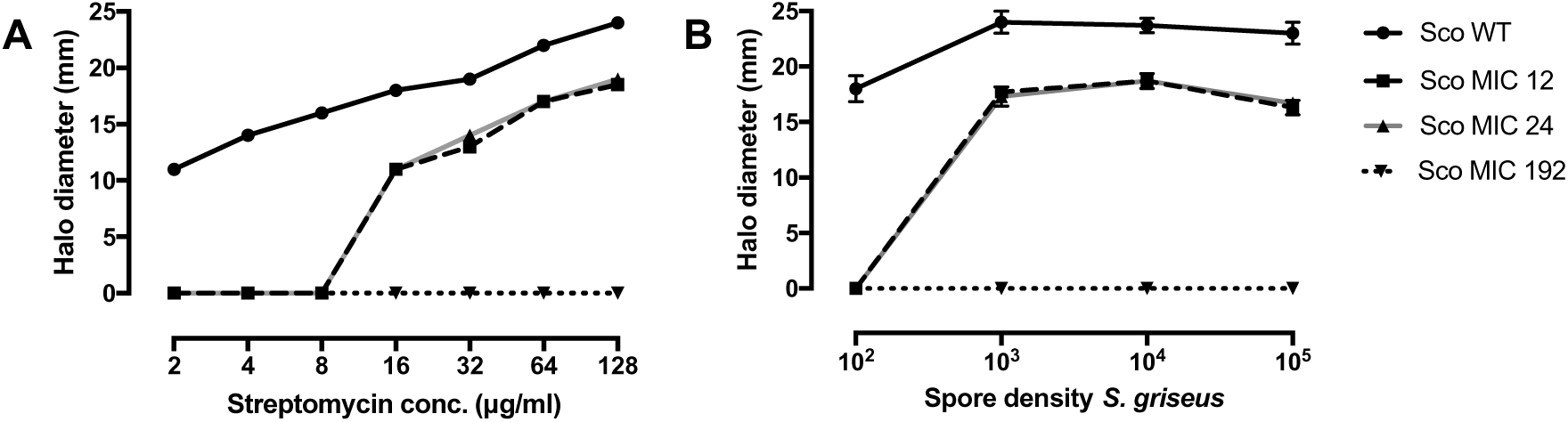
Quantification of *S. griseus* streptomycin production. (A) Halo size of streptomycin susceptible WT and resistant mutants of *S. coelicolor* when exposed to 10 mm agar plugs extracted from plates supplemented with purified streptomycin (final concentration 2 – 128 μg/ml streptomycin) or (B) from four day old plates inoculated with increasing densities of *S. griseus* spores. Error bars represent standard error of the mean.

### Conditions modifying S. griseus invasion

Having established that *S. griseus* produces high concentrations of streptomycin and that this enables *S. griseus* to outcompete *S. coelicolor* when the two strains are equally common, we next sought to identify conditions that influence the invasability of this strain. We focused specifically on *S. griseus* density and frequency as we predicted that these would impact the ability for an antibiotic producing strain to invade when rare, and on the role of spatial structure. The latter parameter has been shown to be particularly important for bacteriocin invasion by allowing producing strains to benefit locally from their own toxin production by creating competition-free space surrounding the producing colony (Chao and Levin, 1981). Competition experiments in Fig 3A clarify that the fitness of *S. griseus* is strongly density-dependent (One-way ANOVA, F_3,8_ = 58.62, p < 0.0001) and that this species can invade from a minimum of ~ 10^3^ spores/plate when competed against a *S. coelicolor* strain with a streptomycin MIC of 48 ug/ml. We can explain these results in two ways: first, less streptomycin is produced when there are fewer *S. griseus* cells on the plate, thus leading to less inhibition of *S. coelicolor*. Second, when fewer spores are plated, the distance between colonies increases. This means that in the vincinity of a producer there are fewer susceptible colonies to inhibit, which reduces the benefit of producing the antibiotic. To distinguish between these two possibilities we next varied the frequency of *S. griseus* in the population (Fig. 3B), while holding the initial density constant at 10^5^ spores per plate. This enabled us to vary the amount of streptomycin produced, while the distance to neighbouring *S. coelicolor* colonies remained constant. The results of these experiments reveal that the fitness benefits of antibiotic production by *S. griseus* are significantly frequency dependent (one-way ANOVA, F_2, 15_ = 71.39, p < 0.0001), and increase with their relative frequency (Fig. 3B). However, these experiments also indicate that the threshold for invasion is relatively high, and that *S. griseus* needs to reach at least 1% of the population before it benefits from antibiotic production. This result raised the question of how *S. griseus* could reach a frequency of 1% from much lower initial values, or if there are conditions that could lower this threshold to permit invasion from fewer cells.

**Fig. 3.**
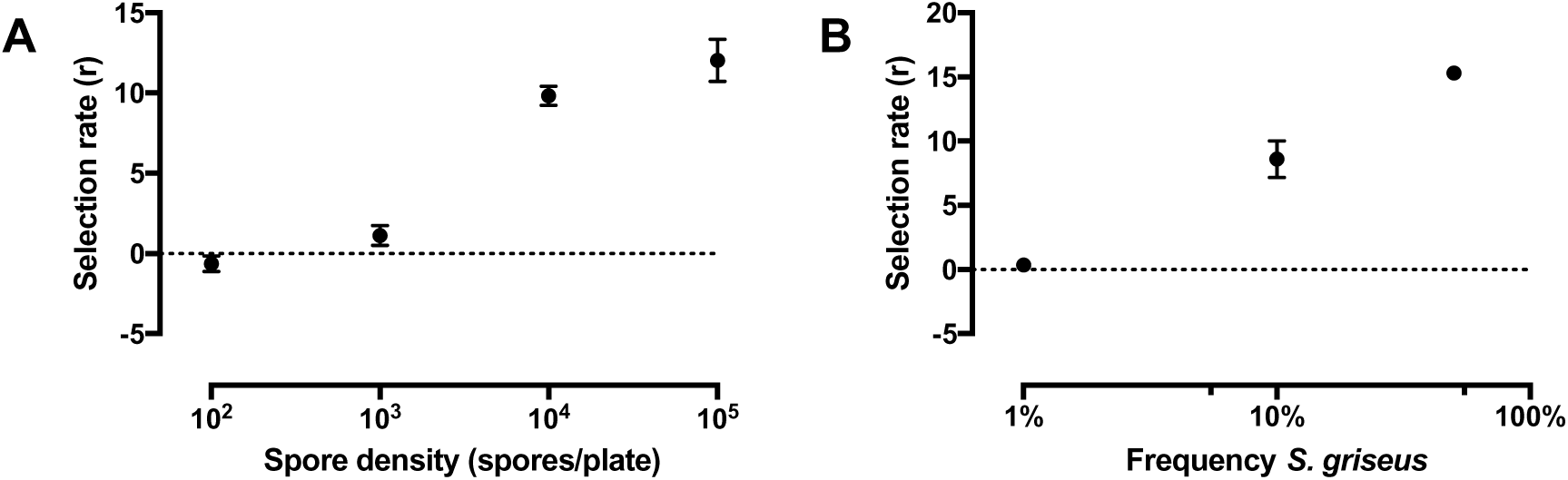
Competition between *S. griseus* and *S. coelicolor* at different densities or frequencies. (A) Fitness of *S. griseus* competing with an intermediate resistant *S. coelicolor* (MIC 48 μg/ml) at equal frequencies but varying spore densities (10^2^ – 10^5^) on the plate. (B) Fitness of *S. griseus* competing against *S. coelicolor* WT at a spore density of 10^5^ spores/plate from different frequencies (1%, 10% and 50%) in the population. Error bars represent standard error of the mean.

To address this issue, we set up experiments to determine the invasion threshold of *S. griseus* in conditions where spatial structure was either maintained over the course of eight serial passages, or was periodically destroyed. These experiments were in part motivated by our observations that we could observe very small inhibition zones surrounding *S. griseus* colonies when competing with *S. coelicolor*, even when *S. griseus* was at low frequencies (see Fig. 4A inset). Although these tiny inhibition zones were insufficient to provide short-term fitness benefits (Fig 3B), we hypothesized that they might permit *S. griseus* to expand from these regions if local structure were maintained. At each transfer, we used replica plating to maintain the spatial structure in each replicate across time. To destroy spatial structure we simply rotated the plate for the next transfer cycle onto the velvet used for replica plating. Consistent with our predictions, we found that maintaining spatial structure enabled invasion from much lower frequencies. When retaining the spatial structure, *S. griseus* was able to invade from an initial frequency of as low as 0.001% and then become fixed in the population (Fig. 4A). Given the initial total inoculation density of 10^5^ spores, this indicates that no more than 10 spores are required for invasion. By contrast, when the spatial structure was destroyed at each transfer cycle, the threshold for invasion increased ~100 fold to 0.1% (Fig. 4B). The cause of these results is clearly illustrated in Fig. 4C (and more detailed in Fig. S2), showing that minute halos expand through time and then eventually coalesce when spatial structure is maintained, but that these halos disappear when it is destroyed. However, it is important to note that spatial structure is in itself insufficient to enable invasion, because when *S. griseus* is competed against a highly resistant strain of *S. coelicolor*, its invasion is prevented even though it was inoculated at a relatively high frequency (Fig 4D). These results further support the conclusion that streptomycin production and sensitivity are the key factors driving the population dynamics of these two species.

**Fig. 4.**
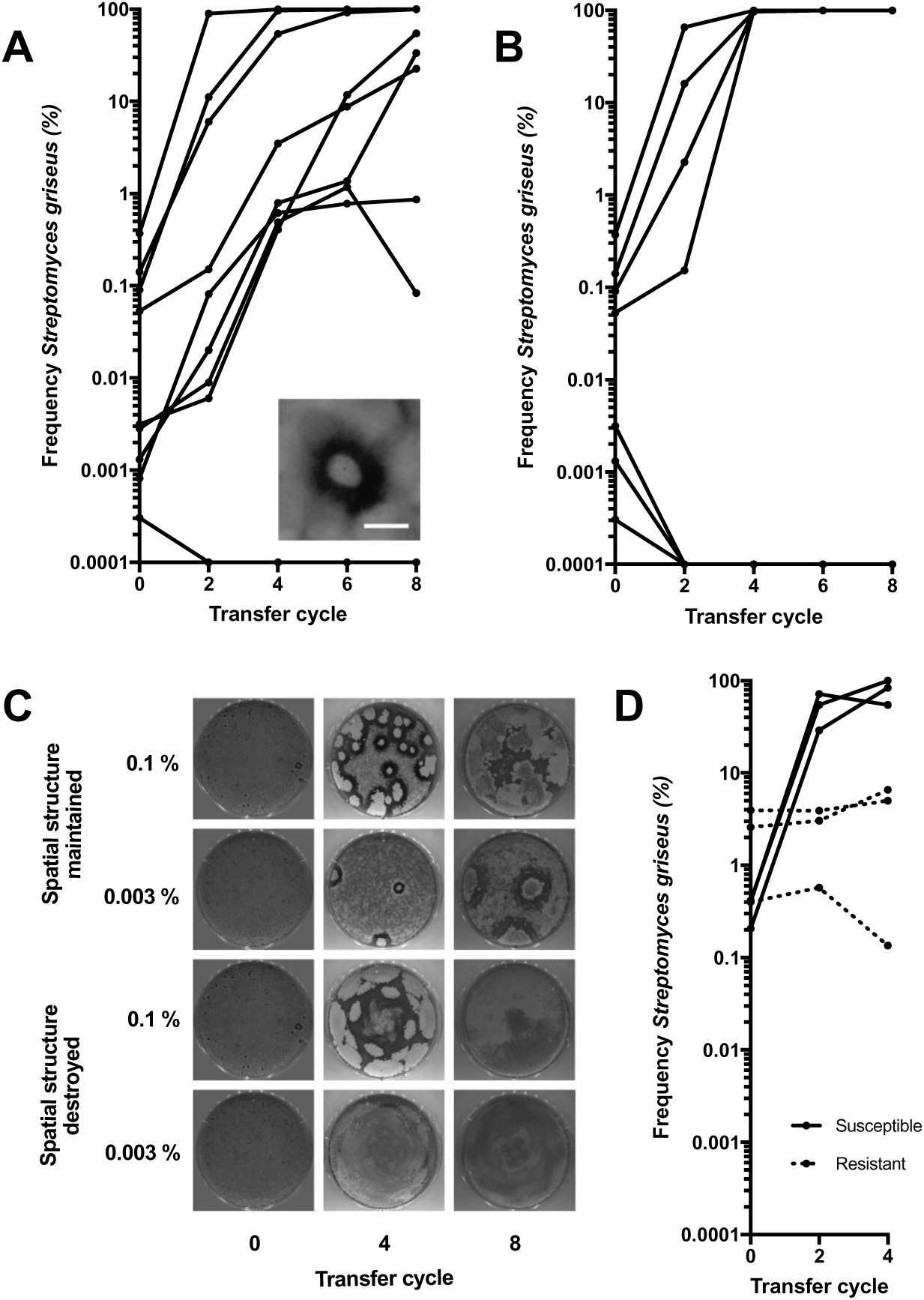
Changes in frequency of *S. griseus* in competition with streptomycin susceptible *S. coelicolor* WT when spatial structure is (A) maintained by replica plating or (B) destroyed by rotating the plate on the velvet used for replica plating in different directions. Inset in A shows a representative image of an inhbition zone around a *S. griseus* colony, scale bar 500 μm. (C) Representative images of plates from A and B at different initial *S. griseus* frequencies. (D) Frequency of *S. griseus* in competition with streptomycin susceptible or highly resistant *S. coelicolor* (MIC 192 ug/ml) when spatial structure is maintained.

## Discussion

Antibiotic production by microbes is ubiquitous in nature, with streptomycin being one of the most commonly produced antibiotics; it has been estimated that 1% of randomly screened actinomycetes from around the globe can synthesize this antibiotic (Baltz, 2008). Although this suggests that antibiotic production confers benefits, very little is known about the population dynamics of antibiotic production and the conditions that influence how antibiotic producers become established. Here we focus on the importance of streptomycin for its producer *S. griseus* during competitive interactions with susceptible and resistant strains of *S. coelicolor*. Our results show that antibiotic production occurs at high levels and that this enables *S. griseus* to kill and therefore invade a population of drug-susceptible competitors. However, this only occurs if *S. griseus* is numerous and at fairly high frequencies, otherwise it fails to outcompete *S. coelicolor*.

These results from short-term competition experiments closely mirror those seen for colicin production in *E. coli*, where in mass action enviroments, producing cells must be moderately common to benefit from colicin production. This leads to two altnernative outcomes; colicin producers fix or they go extinct. As classically shown by Chao and Levin, the reason for this result is that the costs of colicin production, including cell lysis necessary for secretion, exceed the benefits of production in an environment with high rates of diffusion (Chao and Levin, 1981). Although antibiotic secretion is not lethal in streptomycetes, it is expected to be metabolically expensive, with pathways for secondary metabolites comprising ~5% of Streptomyces genomes (Challis and Hopwood, 2003; Nett *et al.*, 2009). Streptomycin production requires the growth-dependent accumulation of A-factor, a small signalling molecule, which through the pathway-specific regulator strR results in the transcription of the streptomycin cluster, a 31 kb gene cluster consisting of 27 genes, as well as the formation of aerial hyphae (Ohnishi *et al.*, 2008; Distler *et al.*, 1992; Bibb, 2005). Similarly to rare colicin producers, when *S. griseus* is rare, it produces insufficient streptomycin in the competition environment to invade, a result that may be partly due to a failure to activate streptomycin production via accumulated A-factor. Even though we were unable to detect a fitness benefit of streptomycin production under these conditions, we often observed extremely small zones of clearance around minute *S. griseus* colonies. This suggests the possibility that continued cultivation would allow *S. griseus* to thrive, as long as cells could directly benefit from their own antibiotic production. Serial transfer experiments where the spatial structure was either retained or periodically destroyed indeed revealed that spatial structure lowered the threshold of invasion by more than 100-fold to fewer than 10 total cells, in parallel with the classic results from Chao and Levin (1981).

The temporal dyanamics of these experiments (Fig. 4C and S2), help to explain our results. Rare streptomycin producers secrete sufficient amounts of the antibiotic to create a small zone of inhibition, thereby freeing space and resources for their growth. With continued cultivation over several transfer cycles, these halos continue to expand radially as the size of the producing colony of *S. griseus* grows. Eventually, halos from separate colonies coalesce, leading to rapid fixation of the antibiotic-producing strain. Because the benefits of antibiotic production remain local, the costs of production can be overcome. However, when spatial structure is destroyed, these local benefits are diluted. Zones of inhibition never expand, because cell densities of founding colonies remain low, so the benefits of streptomycin production remain unchanged through time.

Streptomycetes in soil live on soil grains where overall cell densities are anticipated to be heterogenous and patchy (Probandt *et al.*, 2018). One of the challenges to understanding the role of antibiotics in nature is that their estimated concentrations in bulk soil are extremely small. Aside from technological limitations that may contribute to these estimates, our results show that as long as competition remains local, low antibiotic concentrations in bulk soil are not informative of their potential benefits because producing cells can still inhibit and kill local competitors. Additionally, we and others have shown that even if drug concentrations are low (up to 100-fold lower than the MIC), they are sufficient to rapidly select for antibiotic resistance (Gullberg *et al.*, 2011; Westhoff *et al.*, 2017). Thus the coexistence, at small spatial scales, of bacterial strains that produce and are resistant to antibiotics is most consistent with the argument that antibiotics are used to mediate competitive interactions.

It is important to note, however, that our experiments differ in many ways from the conditions bacteria face in nature. First, rates of diffusion and the availability of resources will markedly differ in highly heterogenous soil compared to a homogeneous agar plate, where antibiotic diffusion is essentially unconstrained and resources are high. Second, while cells are uniformly distributed on our agar plates, they will be more patchily distributed in soil. Microscopic analysis of sand grains from marine sediments revealed that colonization is highly uneven, with protected areas on individual grains being more densely populated than exposed areas (Probandt et al., 2018). Because the fitness of *S. griseus* scales with density, the colonization density in nature will undoubtedly influence the benefits of antibiotic production. Finally, our experiments do not include longer-term interactions that include the evolution of antibiotic resistance. It is possible that *de-novo* resistant strains would exclude antibiotic producers. However, under local competition, these strains could also facilitate the coexistence of antibiotic production, resistance and susceptibility, as has been observed in the real-life rock-paper-scissor dynamics of colicins, both *in vitro* and *in vivo* (Kerr *et al.*, 2002; Kirkup and Riley, 2004). Considering these factors under more environmentally realistic conditions is an obvious and important next step in our work. To summarize, our results indicate that antibiotics can be used as offensive weapons to invade established populations of competitors from a low frequency and suggest that structured habitats are favourable for this invasion and thereby for the evolution of antibiotic producers. They also suggest that bulk-soil estimates of antibiotic concentrations may be misleading with respect to the role of these compounds in nature, and instead argue for the importance of estimating drug concentrations at small spatial scales that better reflect the competitive arena where these metabolites are used.

## Materials and Methods

### Strains and culturing conditions

Two Streptomyces species were used in this study: the streptomycin producing *Streptomyces griseus* IFO13350 (MIC 92 μg/ml) and the streptomycin sensitive *Streptomyces coelicolor* A(3)2 M145 (MIC 2 μg/ml) carrying an integrated pSET152 plasmid conferring apramycin resistance. Spontaneous streptomycin resistant mutants of *S. coelicolor* A(3)2 M145 described in Westhoff et al. (2017) were also used. Briefly, to obtain these mutants, spores were plated on antibiotic concentrations above the MIC and resistant colonies were picked after several days and tested for MIC. We selected low level resistant (MIC 12, 24 and 48 μg/ml respectively) and high level resistant (MIC 192 μg/ml) strains for these experiments. Strains were transformed with the integrating pSET152 plasmid conferring apramycin resistance for the competition experiments.

Strains were grown routinely for four days at 30 °C on Soy Flour Mannitol Agar (SFM) containing 20 g Soy Flour (Biofresh Belgium, Onze-Lieve-Vrouw-Waver, Belgium), 20 g Mannitol (Merck KGaA, Damstadt, Germany) and 15 g agar (Hispanagar, Burgos, Spain) per liter (pH 7.2-7.4). High-density spore stocks were generated by uniformly spreading plates with 50 μl of spore containing solution. After several days of growth, spores were harvested with a cotton disc soaked in 3 ml 30% glycerol after which spores were extracted from the cotton by passing the liquid though an 18g syringe to remove the vegetative mycelium. Spore stocks were titred and stored at –20 °C.

### Competition experiments

We carried out competition experiments between *S. griseus* and the streptomycin susceptible WT or resistant mutants using streptomycin and apramycin resistance as markers. Competition experiments were initiated by mixing strains at the given frequencies and plating 10^5^ spores unless otherwise indicated. To determine the fraction of our innoculum that was streptomycin or apramycin resistant, we simultaneously plated a dilution of this mix on SFM containing 40 μg/ml streptomycin sulphate (Sigma, St. Louis, MO, USA) or 50 μg/ml apramycin sulphate (Duchefa Biochemie, Haarlem, The Netherlands). After 4 days of growth the plates were harvested and the number of each competitor quantified following plating on SFM with streptomycin or apramycin. Following Travisano and Lenski (1996), the selection rate (r) was calculated as the difference in the Malthusian parameters of both strains: r = ln[*S. griseus*(*t*=4)/*S. griseus*(*t*=0)]-(ln[*S. coelicolor*(*t*=4)/*S. coelicolor*(*t*=0)]), where *t* is the time in days of growth after inoculation.

### Quantifying streptomycin production

We developed a halo assay to quantify the production of streptomycin by *S. griseus*. We prepared plates with known concentrations of streptomycin and plates with varying inoculation densities of *S. griseus* spores and incubated these for 4 days at 30 °C. We took 10 mm agar plugs from these plates using the back end of a sterile 1 ml pipette tipe and removed the top 2 mm to remove the *S. griseus* mycelium. We incubated these plugs for 3 days on 50 μg/ml apramycin SFM plates (to prevent any residual *S. griseus* growth) inoculated with 10^5^ spores of streptomycin susceptible or resistant *S. coelicolor* before we measured the halo diameter.

### Serial transfer experiments

To determine the effect of spatial structure on invasion, *S. coelicolor* and *S. griseus* were mixed at the indicated frequencies and 10^5^ spores were plated and grown for 4 days. An imprint of this plate (resulting in transfer 0) was made on a velveteen cloth and two plates were replicated from this: 1) a plate was replicated in the same orientation to maintain the spatial orientation of the original plate; and 2) a plate was replicated by pressing the plate on the velveteen cloth in different orientations to destroy the spatial orientation. From this point plates continued to be replicated in the same manner after 4 days of growth for a total of 4 or 8 transfers depending on the experiment. Before each transfer the plates were imaged using a flat bed scanner. After every transfer the remaining spores on the plate were harvested and the ratio of *S. griseus* and *S. coelicolor* was quantified following plating on SFM with streptomycin or apramycin.

## Supporting information

## Acknowledgements

This work was financially supported by a grant from the Dutch National Science Foundation (NWO) to DER.

## References

Abrudan MI, Smakman F, Grimbergen AJ, Westhoff S, Miller EL, van Wezel GP, et al. (2015). Socially mediated induction and suppression of antibiosis during bacterial coexistence. Proc Natl Acad Sci U S A 112: 11054–11059.

Baltz RH. (2008). Renaissance in antibacterial discovery from actinomycetes. Curr Opin Pharmacol 8: 557–563.

Barka EA, Vatsa P, Sanchez L, Gaveau-vaillant N, Jacquard C, Klenk H-P, et al. (2016). Taxonomy, Physiology, and Natural Products of Actinobacteria. Microbiol Mol Biol Rev 80: 1–43.

Bérdy J. (2005). Bioactive Microbial Metabolites. J Antibiot (Tokyo) 58: 1–26.

Bibb MJ. (2005). Regulation of secondary metabolism in streptomycetes. Curr Opin Microbiol 8: 208–215.

Challis GL, Hopwood DA. (2003). Synergy and contingency as driving forces for the evolution of multiple secondary metabolite production by Streptomyces species. Proc Natl Acad Sci 100: 14555–14561.

Chao L, Levin BR. (1981). Structured habitats and the evolution of anticompetitor toxins in bacteria. Proc Natl Acad Sci 78: 6324–6328.

D’Costa VM, McGrann KM, Hughes DW, Wright GD. (2006). Sampling the Antibiotic Resistome. Science (80-) 311: 374–377.

Davies J. (2006). Are antibiotics naturally antibiotics? J Ind Microbiol Biotechnol 33: 496–499.

Davies J, Spiegelman GB, Yim G. (2006). The world of subinhibitory antibiotic concentrations. Curr Opin Microbiol 9: 445–453.

Distler J, Mansouri K, Mayer G, Stockmann M, Piepersberg W. (1992). Streptomycin biosynthesis and its regulation in Streptomycetes. Gene 115: 105–111.

Ghoul M, Mitri S. (2016). The Ecology and Evolution of Microbial Competition. Trends Microbiol 24: 833–845.

Gullberg E, Cao S, Berg OG, Ilbäck C, Sandegren L, Hughes D, et al. (2011). Selection of resistant bacteria at very low antibiotic concentrations. PLoS Pathog 7: e1002158.

Hibbing ME, Fuqua C, Parsek MR, Peterson SB. (2010). Bacterial competition: surviving and thriving in the microbial jungle. Nat Rev Microbiol 8: 15–25.

Hopwood DA. (2007). Streptomyces in nature and medicine: the antibiotic makers. Oxford University Press: New York.

Kerr B, Riley MA, Feldman MW, Bohannan BJM. (2002). Local dispersal promotes biodiversity in a real-life game of rock-paper-scissors. Nature 418: 171–174.

Kirkup BC, Riley MA. (2004). Antibiotic-meidated antagonism leads to a bacterial game of rock-paper-scissors in vivo. Nature 428: 694–696.

Laskaris P, Tolba S, Calvo-Bado L, Wellington L. (2010). Coevolution of antibiotic production and counter-resistance in soil bacteria. Environ Microbiol 12: 783–796.

Martínez JL. (2008). Antibiotics and Antibiotic Resistance. Science (80-) 321: 365–367.

Nett M, Ikeda H, Moore BS. (2009). Genomic basis for natural product biosynthetic diversity in the actinomycetes. Nat Prod Rep 26: 1362–1384.

Ohnishi Y, Ishikawa J, Hara H, Suzuki H, Ikenoya M, Ikeda H, et al. (2008). Genome sequence of the streptomycin-producing microorganism Streptomyces griseus IFO 13350. J Bacteriol 190: 4050–4060.

Pfuetze KH, Pyle MM, Hinshaw HC, Feldman WH. (1955). The first clinical trial of streptomycin in human tuberculosis. Am Rev Tuberc 71: 752–4.

Probandt D, Eickhorst T, Ellrott A, Amann R, Knittel K. (2018). Microbial life on a sand grain: From bulk sediment to single grains. ISME J 12: 623–633.

Raaijmakers JM, Mazzola M. (2012). Diversity and Natural Functions of Antibiotics Produced by Beneficial and Plant Pathogenic Bacteria. Annu Rev Phytopathol 50: 403–424.

Schatz A, Bugie E, Waksman SA. (1944). Streptomycin, a Substance Exhibiting Antibiotic Activity against Gram-Positive and Gram-Negative Bacteria. Proc Soc Exp Biol Med 55: 66–69.

Westhoff S, van Leeuwe TM, Qachach O, Zhang Z, van Wezel GP, Rozen DE. (2017). The evolution of no-cost resistance at sub-MIC concentrations of streptomycin in Streptomyces coelicolor. ISME J.

Wiener P. (2000). Antibiotic production in a spatially structured environment. Ecol Lett 3: 122–130.

Wiener P. (1996). Experimental studies on the ecological role of antibiotic production in bacteria. Evol Ecol 10: 405–421.

Yim G, Wang HH, Davies J. (2006). The truth about antibiotics. Int J Med Microbiol 296: 163–170.

