## Supplementary material for "Spatial structure increases the benefits of antibiotic production in *Streptomyces griseus*"

### Supplemental Information

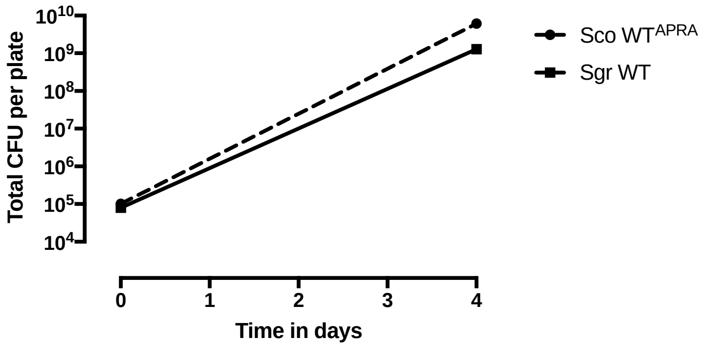


Figure S1. Total number of spores (CFU) initially and after four days of growth of *S. coelicolor* WT and *S. griseus* grown seperately on SFM.

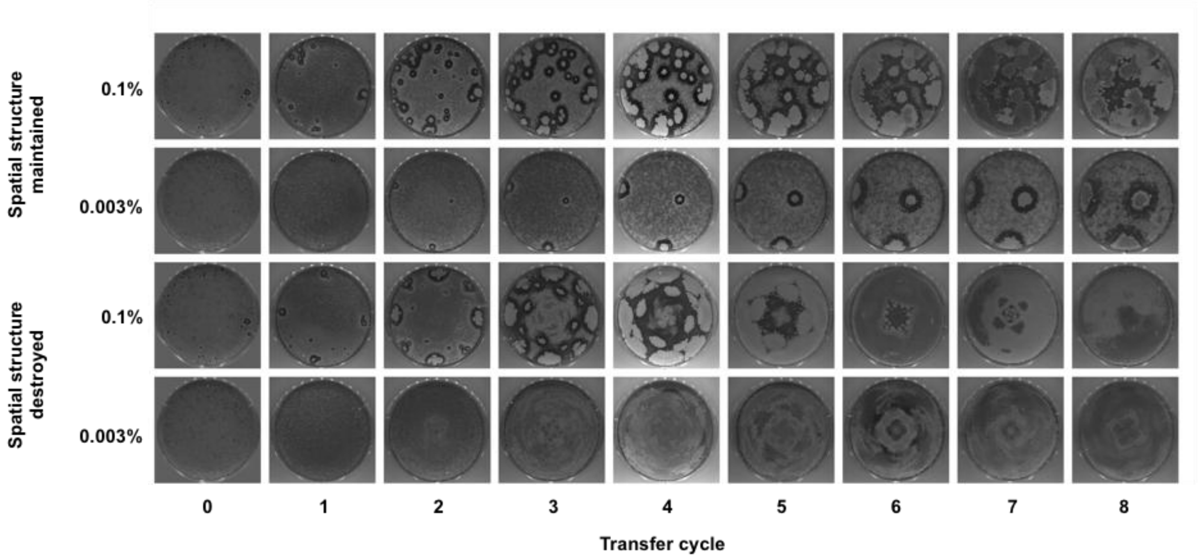


Fig. S2. Representative images of plates from Fig. 4A and B at different initial *S. griseus* frequencies at every transfer cycle.
